# Engineered receptor binding domain immunogens elicit pan-sarbecovirus neutralizing antibodies outside the receptor binding motif

**DOI:** 10.1101/2020.12.07.415216

**Authors:** Blake M. Hauser, Maya Sangesland, Evan C. Lam, Kerri J. St. Denis, Jared Feldman, Ashraf S. Yousif, Timothy M. Caradonna, Ty Kannegieter, Alejandro B. Balazs, Daniel Lingwood, Aaron G. Schmidt

## Abstract

Effective countermeasures are needed against emerging coronaviruses of pandemic potential, similar to severe acute respiratory syndrome coronavirus 2 (SARS-CoV-2). Designing immunogens that elicit broadly neutralizing antibodies to conserved viral epitopes on the major surface glycoprotein, spike, such as the receptor binding domain (RBD) is one potential approach. Here, we report the generation of homotrimeric RBD immunogens from different sarbecoviruses using a stabilized, immune-silent trimerization tag. In mice, we find that a cocktail of these homotrimeric sarbecovirus RBDs elicits antibodies to conserved viral epitopes outside of the ACE2 receptor binding motif (RBM). Importantly, these responses neutralize all sarbecovirus components even in context of prior SARS-CoV-2 imprinting. We further show that a substantial fraction of the neutralizing antibodies elicited after vaccination in humans also engages non-RBM epitopes on the RBD. Collectively, our results suggest a strategy for eliciting broadly neutralizing responses leading to a pan-sarbecovirus vaccine.

**Author summary:** Immunity to SARS-CoV-2 in the human population will be widespread due to natural infection and vaccination. However, another novel coronavirus will likely emerge in the future and may cause a subsequent pandemic. Humoral responses induced by SARS-CoV-2 infection and vaccination provide limited protection against even closely related coronaviruses. We show immunization with a cocktail of trimeric coronavirus receptor binding domains induces a neutralizing antibody response that is broadened to related coronaviruses with pandemic potential. Importantly, this broadening occurs in context of an initial imprinted SARS-CoV-2 spike immunization showing that preexisting immunity can be expanded to recognize other related coronaviruses. Our immunogens focused the serum antibody response to conserved epitopes on the receptor binding domain outside of the ACE2 receptor binding motif; this contrasts with current SARS-CoV-2 therapeutic antibodies, which predominantly target the receptor binding motif.

## Introduction

The emergence of the novel severe acute respiratory syndrome coronavirus 2 (SARS-CoV-2) and the subsequent global pandemic has highlighted the disruptive threat posed by viruses for which humans have no prior immunity. Rapid development of potential vaccines has led to an unprecedented 40 candidates already in Phase 3 clinical trials or approved since January 2020; while differing in modality (e.g., mRNA, adenovirus), the primary immunogen for many of these candidates is the SARS-CoV-2 spike ectodomain (1). With the continued global spread of SARS-CoV-2, in conjunction with potential vaccinations, it is likely that a large proportion of the global population will eventually develop an immune response to SARS-CoV-2. However, even after potentially achieving herd immunity sufficient to slow the spread of SARS-CoV-2, there remains a constant threat of emerging coronaviruses with pandemic potential. Indeed, surveillance efforts have identified numerous unique coronaviruses within various animal reservoirs, raising the possibility of zoonotic transmission (2, 3). Such events are likely to increase in frequency as a result of human impact on the environment (4). While the current SARS-CoV-2 pandemic has an estimated infection fatality rate of between ~1-2%, with numerous cases likely undetected, previous SARS-CoV and MERS-CoV outbreaks were more lethal with ~10% and ~35% case fatality rates, respectively, raising the possibility that future novel coronaviruses will potentially have high mortality (5–7). Additionally, elicited immunity to SARS-CoV-2 infection may not protect against even closely related novel coronaviruses from the same sarbecovirus subgenus (8, 9). It is therefore critical to not only address the current pandemic, but also develop vaccine platforms that can be readily adapted to potential emerging coronaviruses.

While we cannot readily predict which coronaviruses will next emerge into the human population, a proactive approach to generate broadly protective immunity is to design immunogens that elicit humoral responses targeting conserved sites on the coronavirus spike glycoprotein. Such sites may include the receptor binding domain which engages host-cell receptors necessary to facilitate viral cell entry; these spike-mediated interactions are conserved across coronaviruses (10). Indeed, a subset of spike-directed antibodies from convalescent patients can potently neutralize SARS-CoV-2; comparable neutralizing antibodies against SARS-CoV and MERS-CoV have also been identified (6, 11–16). Some spike-directed antibodies that neutralize SARS-CoV-2 also bind the SARS-CoV spike protein, highlighting the presence of cross-reactive neutralizing epitopes (11, 17, 18). Multimerized versions of the receptor binding domains (RBDs) of several coronaviruses have previously been shown to be potent immunogens (19). Here, we describe a customizable vaccine that elicits pan-sarbecovirus neutralization against both SARS-CoV-2 and the potentially emergent WIV1-CoV (20). This approach focuses antibody responses to conserved, protective RBD epitopes shared across sarbecoviruses. Its flexible nature allows facile interchanging of potential vaccine strains updated to confer neutralization against new emerging coronaviruses. Responses focused to these conserved epitopes maintain potent SARS-CoV-2 neutralization activity despite minimal ACE2 receptor binding motif coverage. Importantly, this approach was applied in context of “pre-existing” SARS-CoV-2 humoral immunity with the goal of broadening neutralization while simultaneously boosting the neutralizing antibody response to SARS-CoV-2, similar to the “back boost” effect of seasonal influenza immunizations (21). Thus, it has the potential to provide protection against currently circulating SARS-CoV-2, while proactively generating neutralizing antibody responses against emerging coronaviruses.

## Results

### Designing a Trimerized Receptor Binding Domain Construct Using a Non-Immunogenic Tag

We designed a cystine-stabilized and hyperglycosylated variant of a GCN4 trimerization tag to generate a homotrimeric immunogen “cassette” to rapidly exchange RBDs from various coronaviruses (**Fig. 1A**). While the two cysteines are within one subunit, they are engineered to form an intermolecular disulfide with an adjacent subunit when the trimer is formed. Using a hyperglycosylated GCN4 allows the RBDs to remain trimerized while the tag is “immune silent” (22). As a proof-of-concept for our immunization approach, we selected the SARS-CoV, SARS-CoV-2, and WIV1-CoV RBDs as our starting immunogens. We overexpressed RBD homotrimers in mammalian cells (Expi293F cells) to maximize glycan complexity and purified to homogeneity via immobilized metal affinity chromatography followed by size exclusion chromatography; the trimeric species was confirmed using SDS-PAGE analysis under non-reducing conditions (**Fig. 1B-C**). Their antigenicity was assayed using conformational-specific antibodies CR3022 and/or B38 using biolayer interferometry; the RBD homotrimers had comparable affinities in comparison to RBD monomers (**Fig. S1**). The 8xHis purification tags were removed by HRV 3C protease enzymatic cleavage prior to immunization.

**Fig. 1.**
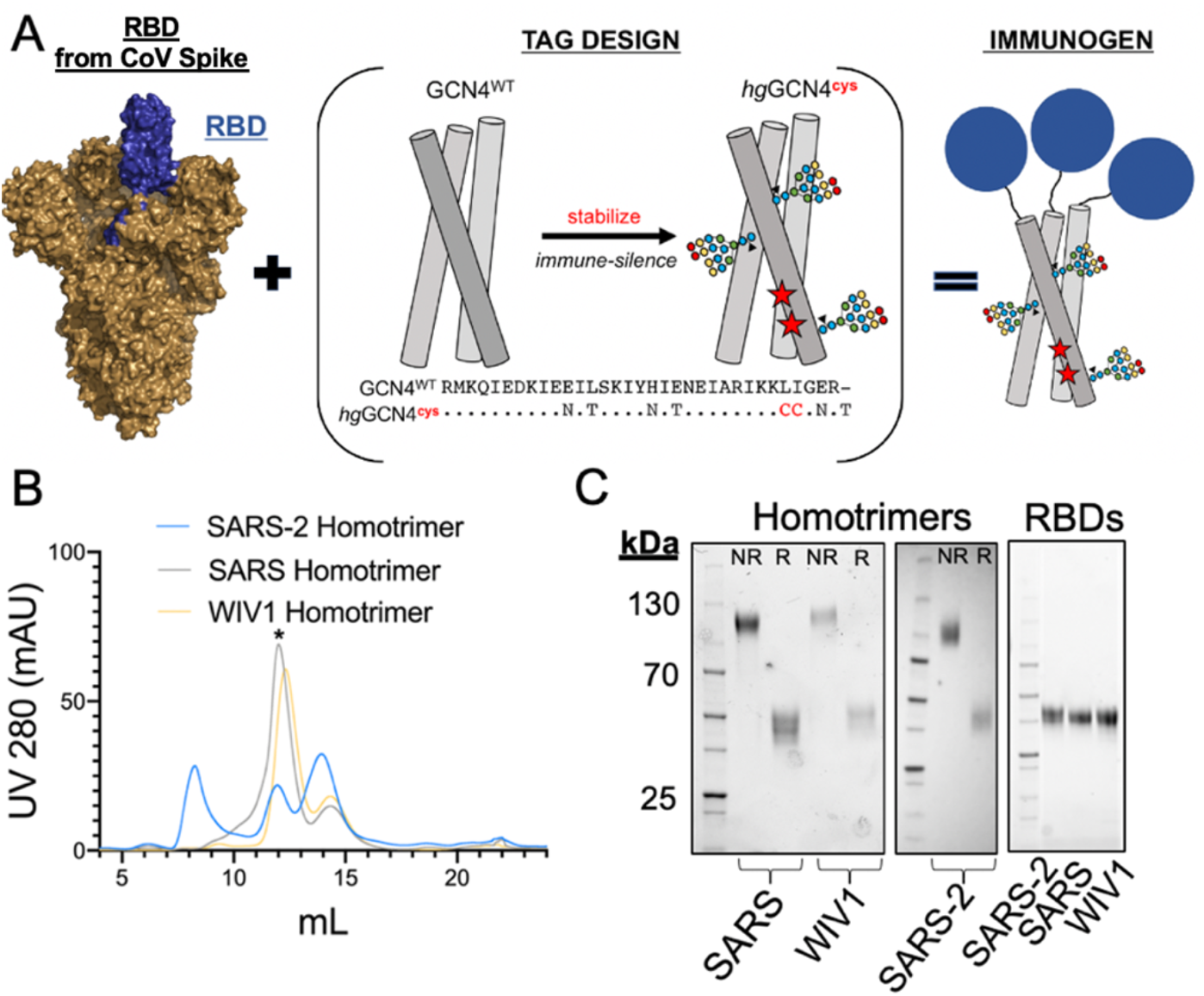
Immunogen design, expression, and purification. (**A**) Design schematic for generating RBD homotrimers appended onto a cystine-stabilized (red stars) hyperglycosylated GCN4 tag. (PDB: 6VSB) (**B**) Representative size exclusion trace with (*) marking the homotrimeric construct. Fractions in this peak were pooled and used for immunizations. **(C)** SDS-PAGE analysis of purified homotrimers following removal of the affinity purification tags under non-reducing (NR) and reducing (R) conditions. The engineered disulfide bond at the C-terminus of the hyperglycosylated GCN4 tag separated under reducing conditions. Panel includes monomeric RBDs run under reducing conditions for comparison. (Related to **Fig. S1**)

### Immunization Regimens Generate Cross-Reactive Antibody Responses

To understand how preexisting immunity to SARS-CoV-2, whether acquired through natural infection or vaccination, could affect immune responses to our immunogens, we primed our cohorts with recombinant spike or RBD protein (23, 24). Our two control arms followed this prime with subsequent homologous boosts of recombinant SARS-CoV-2 spike (“Spike” cohort) or RBD (“RBD” cohort); the latter is necessary for comparing the effect on immunogencity of monomeric versus trimeric RBDs. Our experimental arms both received the spike prime and included a SARS-CoV-2 RBD homotrimer boost (“Homotrimer” cohort) and an equimolar boosting cocktail of SARS-CoV-2, SARS-CoV, and WIV1-CoV RBD homotrimers (“Cocktail” cohort) (**Fig. 2A**). All cohorts received the same total amount of protein in each immunization, 20 μg, adjuvanted with Sigma Adjuvant (25). Immunizations were performed at days 0, 21, and 42. We evaluated the serum response against coronavirus-derived antigens using ELISAs, including SARS-CoV-2 spike, RBDs from SARS-CoV-2, SARS-CoV, WIV1, as well as a SARS-CoV-2 ΔRBM RBD with two engineered glycans that abrogate ACE2 engagement (**Fig. 2B, S2A, S3A-C**). All four immunization regimens resulted in similar patterns of serum reactivity. Each cohort demonstrated a significant decrease in reactivity against SARS-CoV RBD as compared to the SARS-CoV-2 RBD or spike, though the magnitude of this difference was relatively small. The cohort which received the RBD homotrimer cocktail boost had the highest overall endpoint titers, while the cohort that received the monomeric SARS-CoV-2 RBD had the lowest overall endpoint titers; the latter observation is consistent with other previous reports and likely due to the inefficiency of monomeric RBDs effectively stimulating B cell receptors. We confirmed the non-immunogenic nature of the hyperglycosylated GCN4 tag by assaying sera from the wildtype homotrimer cocktail boost against another viral glycoprotein, hemagglutinin, with the same tag (**Fig. S2B**).

**Fig. 2.**
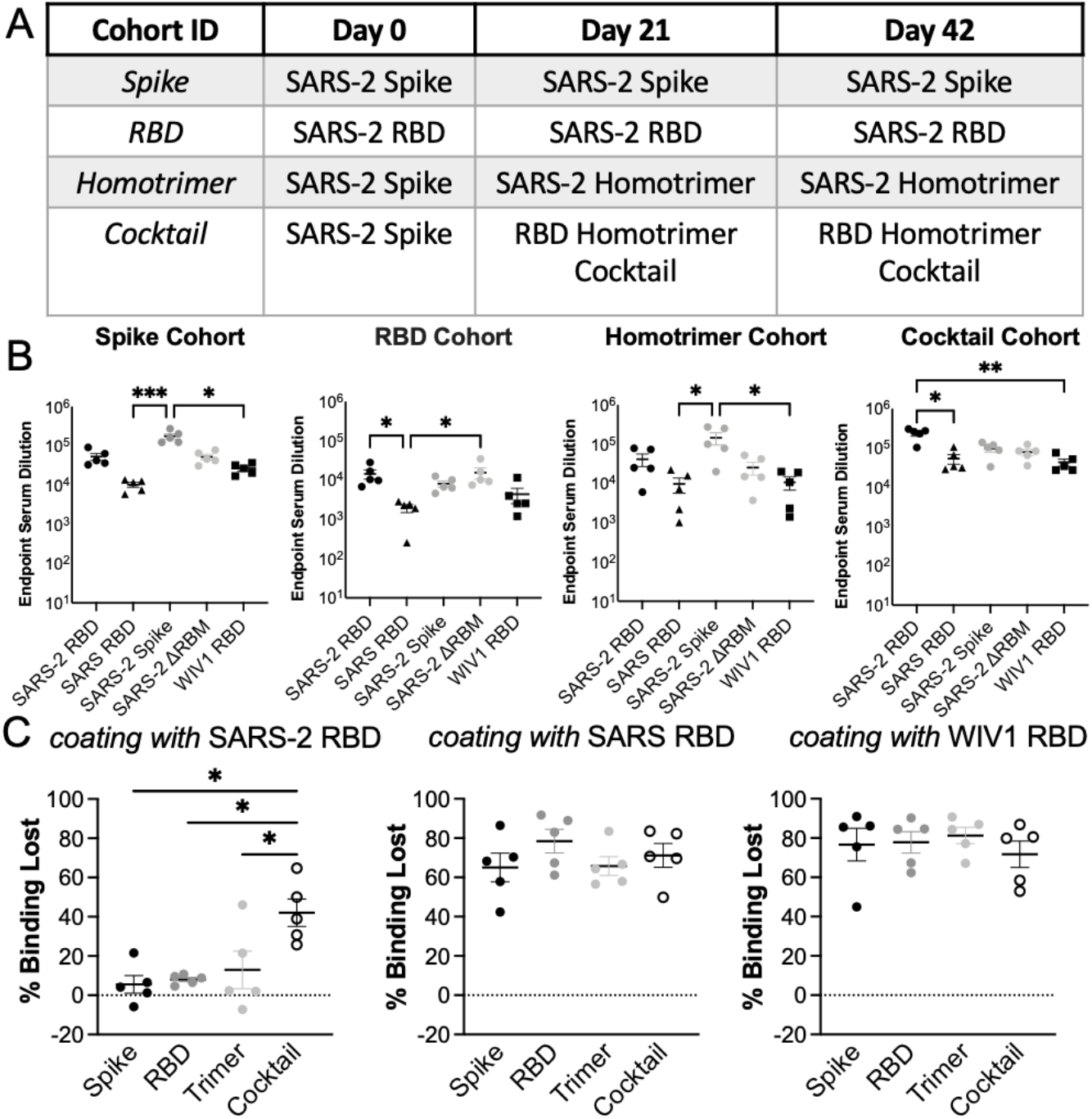
Serum response to immunization regimens. (**A**) Four immunization cohorts were used for this study (n=5 mice, per cohort). (**B**) Serum was assayed in ELISA at day 56 with different coronavirus antigens. (**C**) Percent of binding lost in competition ELISAs using S309 and CR3022 vs. no IgG and SARS-CoV-2, SARS-CoV, and WIV1-CoV RBDs as coating antigens. Statistical significance was determined using Kruskal-Wallis test with post-hoc analysis using Dunn’s test corrected for multiple comparisons (* = p < 0.05, ** = p < 0.01, *** = p < 0.001); ns = not significant. (Related to **Fig. S2**)

### Immunization with a Cocktail of Homotrimeric Receptor Binding Domains Focuses the Immune Response to Cross-Reactive Epitopes

We next evaluated whether the serum response was directed towards cross-reactive, and potentially broadly neutralizing RBD epitopes. Conservation across SARS-CoV-2, SARS-CoV, and WIV1-CoV RBDs primarily occurs outside of the ACE2 receptor binding motif (RBM). Indeed, the previously characterized CR3022 and S309 antibodies have footprints that together cover much of this conserved region, with epitope buried surface area (BSA) of 917 Å^2^ and 795 Å^2^ respectively in comparison to BSA of 869 Å^2^ for ACE2 (17, 18, 26). We performed serum competition by incubating RBD-coated ELISA plates with IgGs B38, P2B-2F6, CR3022, and S309, representing each of the four previously defined “classes” of SARS-CoV-2 RBD epitopes (27) (**Fig. S2C**). We then assessed binding of mouse serum IgG. In all cohorts, competition with both CR3022 and S309 significantly reduced serum titers against the SARS-CoV and WIV1-CoV RBDs (**Fig. 2C, S2C**). However, only the cohort receiving the RBD homotrimer cocktail showed a significant reduction in serum titers against the SARS-CoV-2 RBD in competition with both CR3022 and S309 (p = 0.0340) (**Fig. 2C**). This result suggests a higher degree of focusing to this region elicited specifically by the homotrimer cocktail immunogens.

### Expanded IgG B Cell Populations Target Cross-Reactive Receptor Binding Domain Epitopes

To compare the observed sera responses, we measured the amount of antigen-specific IgG B cells expanded by the Spike, Homotrimer, and Cocktail immunization regimens. Low antigen-specific ELISA titers for the RBD cohort indicated that we were unlikely to be able to robustly quantitate antigen-specific B cells, therefore mice from that cohort were excluded from subsequent analyses. We engineered a SARS-CoV-2 RBD variant that has two additional glycans on the RBM which effectively block ACE2 engagement (**Fig. S3A-C**). This variant allowed us to bin SARS-CoV-2 spike-directed B cells into 3 populations: those that bound RBM epitopes; those that bound the non-RBM epitopes on the RBD; and those that bound the “remainder” of the spike protein (**Fig. S3D-E**). As a subset of SARS-CoV-2 spike-directed B cells, the proportion of B cells specific for the SARS-CoV-2 RBM and non-RBM portion of the RBD were considerably higher in the cohorts receiving the SARS-CoV-2 RBD homotrimer and RBD homotrimer cocktail boosts (p = 0.0070) (**Fig. 3A**). We also binned B cells that bound to both the SARS-CoV-2 spike and either the SARS-CoV RBD or WIV1-CoV RBD (**Fig. 3B**). We found that cross-reactive IgG B cells predominantly targeted epitopes outside the RBM (the spike remainder a result of the prime), mirroring what we observed in sera responses.

**Fig. 3.**
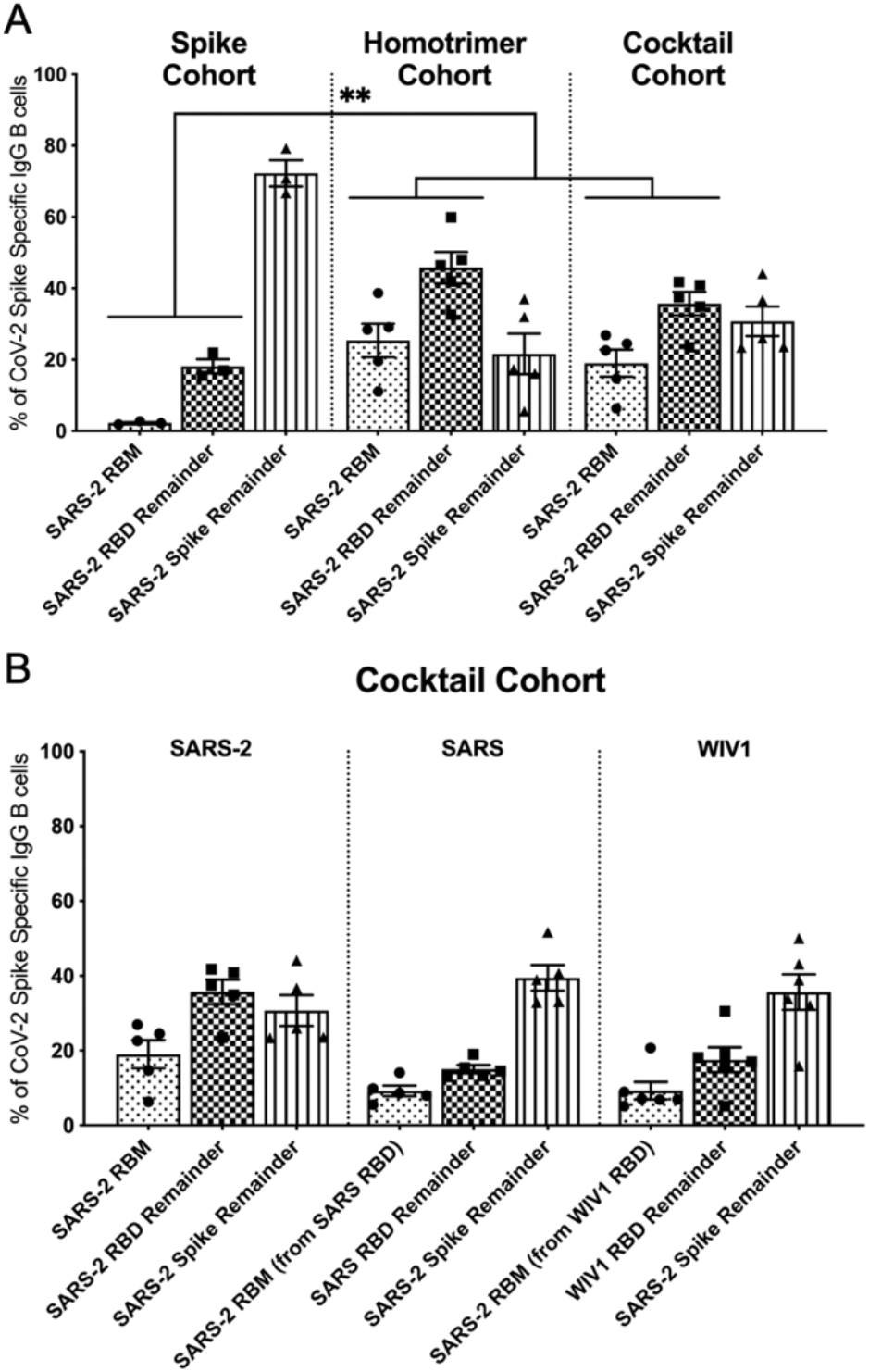
SARS-CoV-2-specific IgG B cells expanded following immunization. **(A)** Spike-directed responses were binned into RBM, RBD remainder (excluding RBM epitopes), and spike remainder (excluding RBD and RBM epitopes) populations using relevant probes in flow cytometry. Data is shown as a percentage of total spike-specific IgG B-cells. We evaluated the distribution of SARS-CoV-2 spike-directed responses and found evidence of focusing towards non-spike RBD epitopes in the cohorts boosted with the SARS-CoV-2 RBD homotrimer and the RBD homotrimer cocktail versus the cohort that received three SARS-CoV-2 spike immunizations (p = 0.0070 via Mann-Whitney U test) (A). (**B**) The cocktail cohort was additionally assayed for SARS-CoV or WIV1-CoV RBD reactivity, as described in (**A**). (Related to **Fig. S3**)

### Elicited Immune Response is Cross-Neutralizing and Targets Non-Receptor Binding Motif Epitopes

We next determined the neutralization potency from each of our cohorts using SARS-CoV-2, SARS-CoV, and WIV1-CoV pseudoviruses (8, 9). We obtained NT50 values when possible; we note that most serum samples for which NT50 values could not be determined still had some weak neutralizing activity (20 out of 24 samples) (**Fig. S4E**). We observed a significant increase in WIV1-CoV neutralization in the cohort that received the homotrimer cocktail boost compared to all other cohorts (p = 0.0012) (**Fig. 4A**). Importantly, this did not result in any significant loss in serum neutralization potency against SARS-CoV-2 (p = 0.6594) (**Fig. 4B**). We also observed higher levels of SARS-CoV neutralization in the cohort that received the homotrimer cocktail boost as compared to the cohort boosted with the SARS-CoV-2 RBD homotrimer and the SARS-CoV-2 monomeric RBD, though this trend was not significant (p = 0.2370) (**Fig. 4C**). The samples from the cohort that received the homotrimer cocktail boost where an NT50 value could not be obtained, nevertheless still show some evidence of weak neutralization.

**Fig. 4.**
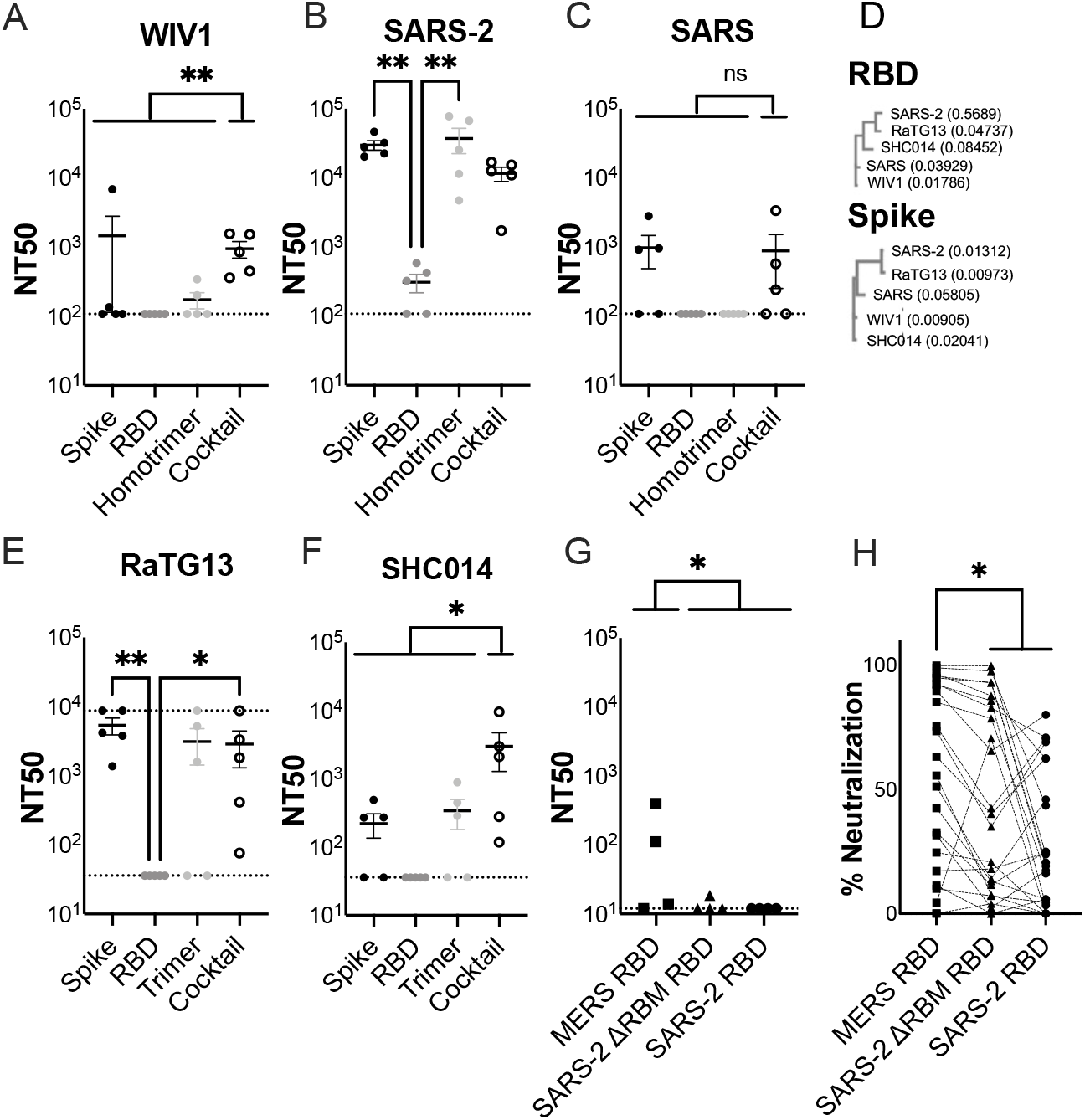
Pan-sarbecovirus serum neutralization using pseudoviruses. Day 56 serum from each cohort was assayed for neutralization against (**A**) WIV1-CoV, (**B**) SARS-CoV-2, (**C**) SARS-CoV, (**E**) RaTG13-CoV, and (**F**) SHC014-CoV pseudoviruses. For the SARS-CoV-2 (**B**) and RaTG13-CoV (**E**) pseudoviruses, statistical significance was determined using Kruskal-Wallis test with post-hoc analysis using Dunn’s test; * = p < 0.05, ** = p < 0.01. There was a significant difference in WIV1-CoV neutralization (Mann-Whitney U test; p = 0.0012) (**A**) and SHC014-CoV neutralization (Mann-Whitney U test; p = 0.0145) (**F**), but there was no significant difference in SARS-CoV neutralization between the control cohorts and the RBD homotrimer cocktail boost cohort (Mann-Whitney U test; p = 0.2370) (**C**). (**D**) Neighbor-joining phylogenetic trees were generated to describe the phylogenetic relationships between the RBDs and spikes of the coronaviruses used in this neutralization panel. Branch lengths are displayed in parentheses. (**G**) Serum adsorption to remove antibodies from pooled sera from each mouse cohort directed towards MERS-CoV RBD (negative control), SARS-CoV-2 ΔRBM RBD, and SARS-CoV-2 RBD. Sera adsorbed with MERS-CoV RBD had significantly greater neutralization than those adsorbed with SARS-CoV-2 ΔRBM RBD and SARS-CoV-2 RBD (Mann-Whitney U test; p = 0.0343). (**H**) Same assay in (**G), except on immune sera from humans who received** the Pfizer (BNT162b2) or Moderna (mRNA-1273) vaccine (Mann-Whitney U test; p = 0.0253). In (**H**), percent neutralization was assessed at a 1:3 dilution of the adsorbed sample. Negative percent neutralization values in (**H**) were set to zero to facilitate analysis. (Related to **Fig. S4**)

To assess cross-neutralization of related coronaviruses that were not included in the RBD homotrimer cocktail cohort, we performed neutralization assays using RaTG13-CoV and SHC014-CoV pseudoviruses, both of which are members of the sarbecovirus subgenus that have been detected in bats but not in humans (**Fig. 4D**) (28, 29). These viruses, along with WIV1, are ACE2-using sarbecoviruses in animal reservoirs that could enter the human population, similar to SARS-CoV and SARS-CoV-2. RaTG13-CoV is closely related to SARS-CoV-2 with 89.6% amino acid identity within the RBD, while SHC014 is more distant with 76.3% identity. For RatG13-CoV, we observed similar neutralization patterns whereby the RBD cohort had significantly lower neutralization than the Spike and Cocktail cohorts. (**Fig. 4D-E**). For SHC014-CoV, however, the cohort that received the RBD homotrimer cocktail boosts showed significantly greater neutralization compared to all other cohorts (p = 0.0145) (**Fig. 4D, F**). Furthermore, the corresponding ELISA titers also show no loss in binding to the SHC014-CoV and RaTG13-CoV RBDs relative to SARS-CoV-2 RBD (**Fig. S2D-E**). Thus, compared to other cohorts, immunization with the RBD homotrimer cocktail resulted in a neutralizing antibody response with both retrospective (e.g., SARS-CoV) and prospective breadth (e.g., WIV1-CoV, SHC014-CoV, and RaTG13-CoV) even in context of preexisting immunity to SARS-CoV-2.

We wanted to determine what fraction of SARS-CoV-2 neutralization could be attributed to the non-RBM RBD-directed response. To that end, we performed adsorption of pooled sera from each cohort using MERS-CoV RBD (negative control), SARS-CoV-2 RBD, and a SARS-CoV-2 RBD with four additional glycans engineered onto the RBM (SARS-CoV-2 ΔRBM RBD) (**Fig. S3B, S4A-D**). This construct was designed to block all RBM-directed antibodies as opposed to the one used in flow cytometry, which only block antibodies that directly compete with ACE2. Samples adsorbed with all three RBD constructs showed decreased neutralization compared to serum due to the dilution involved in the adsorption protocol. Compared to the samples adsorbed using MERS RBD, the samples adsorbed with either the SARS-CoV-2 RBD or the SARS-CoV-2 ΔRBM RBD demonstrated a significant loss of SARS-CoV-2 neutralization (p = 0.0343) (**Fig. 4G, S4E**). This indicates that non-RBM RBD-directed antibodies alone are able to confer significant SARS-CoV-2 neutralization. We performed similar adsorption experiments on 24 post-vaccination human serum samples from a previously characterized vaccine cohort (9). These patients had received either 1 or 2 doses of the Pfizer (BNT162b2) or Moderna (mRNA-1273) vaccines. Similar to our murine observations, human samples adsorbed using either the SARS-CoV-2 RBD or the SARS-CoV-2 ΔRBM RBD lost significant neutralization compared to samples adsorbed using MERS RBD (p = 0.0253) (**Fig. 4H**). Still, samples adsorbed with SARS-CoV-2 ΔRBM RBD did show higher neutralization activity relative to samples adsorbed with SARS-CoV-2 RBD across both the human and mouse samples. This indicates that RBM- and non-RBM antibodies targeting the RBD play important roles in conferring SARS-CoV-2 neutralization.

## Discussion

Here, we demonstrate that a cocktail of homotrimeric sarbecovirus RBDs can effectively generate a neutralizing response to all components and additional related sarbecoviruses without a bias resulting from an initial SARS-CoV-2 imprinting. We find that this cross-neutralizing antibody response is predominantly directed to RBD epitopes outside of the RBM. This contrasts SARS-CoV-2 infection which does not appear to reliably generate cross-neutralizing antibodies (8, 9). Previous analysis into the epitopes targeted by this cross-neutralizing response has not been conducted (30). Furthermore, previously reported vaccine candidates that aim to generate such a cross-neutralizing response have not been shown to successfully focus immunity towards cross-neutralizing epitopes following an initial SARS-CoV-2 spike immunization, but instead were tested only in a naïve homologous prime/boost format (30). However, our results suggest that following an initial SARS-CoV-2 exposure (e.g., vaccination, infection), subsequent boosting with a surveillance-informed selection of sarbecovirus RBD homotrimers could result in pan-sarbecovirus immunity that protects against future pandemics despite targeting non-RBM epitopes. Whereas currently available vaccine candidates, particularly mRNA-based vaccines, appear to largely provide effective protection against currently circulating SARS-CoV-2 variants of concern, it remains unclear whether currently approved vaccines will provide protection against emerging sarbecoviruses (31–35). Immunization with SARS-CoV-2 spike-based vaccines, as well as SARS-CoV-2 infection, results in a significant loss in neutralization against existing pre-emergent sarbecoviruses in both humans and animal models (8, 9, 36). SARS-CoV-2 spike-based vaccines also have reduced *in vivo* protection against SARS-CoV and WIV1 compared to SARS-CoV-2 (36).

Monomeric as well as numerous SARS-CoV-2 multimerized RBD-based vaccine constructs have been published recently and are in various stages of preclinical and clinical testing (19, 30, 37–44). However, these multimerization platforms present additional epitopes that can give rise to a scaffold-specific antibody response, which has the potential to alter hierarchies of immunodominance. Our non-immunogenic hyperglycosylated, cystine-stabilized GCN4 tag improves upon a previous hyperglycosylated version of the tag that showed markedly reduced immunogenicity (22). This may be due to the implementation of cystine-stabilization limiting the accessibility of epitopes within the coiled-coil interface of the GCN4 tag that may not be fully shielded by engineered glycans.

This immunogen cocktail provides a framework upon which further studies to optimize a pan-coronavirus vaccine can build to generate optimal broadly neutralizing antibody responses. This could include antibodies targeting the N-terminal domain as well as the RBD, possibly elicited by vaccine regimens including full-length spike proteins. SARS-CoV-2 pseudovirus neutralization assays may underestimate the contributions of antibodies that target epitopes outside the RBD, though it remains unclear the extent to which this occurs when measuring a polyclonal serum response and against other coronaviruses (45–47). Still, most potently neutralizing monoclonal antibodies and immunization-elicited neutralizing antibodies in humans receiving the approved mRNA vaccines appear to target the RBD, emphasizing the importance of shaping the RBD-directed immune response with any potential future boosting immunizations given the likely impending ubiquity of vaccine-elicited immunity (48–51).

While the durability of vaccine or infection-elicited antibody responses to SARS-CoV-2 remains to be seen, data from seasonal coronaviruses, as well as SARS-CoV and MERS-CoV, suggests that immunity appears to wane after several years and can vary in potency between individuals (52–58). Thus, if herd immunity is not achieved or if antigenic drift of SARS-CoV-2 necessitates reformulation of current vaccines, it may present an opportunity to incorporate immunogens based on emerging coronaviruses identified through surveillance (20); importantly, based on the data presented here, such incorporation would not be at the detriment of neutralizing activity against SARS-CoV-2. Assessing *in vivo* protection efficacy of this vaccine regimen against SARS-CoV-2, in context of preexisting immunity to SARS-CoV-2, is hindered by the observation that a single SARS-CoV-2 spike immunization alone provides notable *in vivo* protection (59).

In addition to informing future immunization regimens, these findings demonstrate that potent cross-neutralizing antibodies can target epitopes outside of the RBM on the SARS-CoV-2 RBD. Our results further demonstrate that non-RBM neutralization via the RBD forms a significant fraction of SARS-CoV-2 immunity elicited in in humans. Many potently neutralizing monoclonal antibodies currently in therapeutic development target RBM epitopes in regions of the RBD with minimal conservation across members of the sarbecovirus subgenus (48, 51). Consequently, these antibodies are unlikely to provide protection against future emerging coronaviruses. Identifying neutralizing antibodies targeting conserved epitopes outside the RBM and the appropriate immunizations strategies and modalities to elicit them may provide the broad protection necessary for pan-coronavirus immunity (11, 18, 49). Furthermore, targeting these non-RBM, but still conserved epitopes, may also reduce the likelihood of antibody escape, which have already been documented for existing SARS-CoV-2 monoclonal antibodies (45, 60). Immunogens that facilitate immune focusing to conserved RBD epitopes, while still presenting the SARS-CoV-2 RBM, can generate a cross-neutralizing antibody response in addition to eliciting SARS-CoV-2-directed antibodies belonging to the potently neutralizing classes of RBM-directed antibodies.

While the currently approved vaccines use an mRNA platform to present SARS-CoV-2 spike, additional immunization strategies use protein-based multivalent constructs (19, 30, 37). Collectively these efforts are designed to elicit both SARS-CoV-2 and, in some cases, pan-coronavirus immunity. The approach described here demonstrates a potential alternate but complementary path to generate neutralizing antibodies against multiple sarbecoviruses even in the context of prior immunity to SARS-CoV-2. Furthermore, this cocktail-based approach using trimeric immunogens parallels the current influenza vaccine composition, which aims to elicit broad immunity to each constituent strain. This proof-of-principle study indicates that a similar surveillance-based approach could be applied to future coronavirus vaccines, that protect against future emerging coronaviruses while also maintaining protection against SARS-CoV-2.

## Materials and Methods

### Receptor Binding Domain and Homotrimer Expression and Purification

Receptor binding domains (RBDs) were designed based on the following sequences: SARS-CoV-2 RBD (Genbank MN975262.1), SARS-CoV RBD (Genbank ABD72970.1), WIV1-CoV RBD (Genbank AGZ48828.1), and MERS-CoV RBD (Genbank AHI48572.1). Constructs were codon optimized by Integrated DNA Technologies, cloned into pVRC, and sequence confirmed by Genewiz. The spike plasmid was obtained from Dr. Jason McLellan at the University of Texas, Austin. It contained a Foldon trimerization domain as well as C-terminal HRV 3C-cleavable 6xHis and 2xStrep II tags. Proteins were expressed in Expi293F cells (ThermoFisher) using Expifectamine transfection reagents according to the manufacturer’s protocols. All proteins included a C-terminal HRV 3C-cleavable 8xHis tag to facilitate purification. Monomeric RBD proteins also contained SBP tags, while homotrimeric constructs contained a previously published hyperglycosylated GCN4 tag with two additional C-terminal cystines (22). A linker with the sequence GAGSSGSG separated each RBD from the hyperglycosylated GCN4 tag. Versions of the MERS-CoV RBD, SARS-CoV-2 ΔRBM RBD with four additional putative glycosylation sites (Figure S5), and SARS-CoV-2 RBD with C-terminal 8xHis and SNAP tags (61) were also generated.

Transfections were harvested after 5 days and clarified via centrifugation. Cell supernatants were passaged over Cobalt-TALON resin (Takara) for immobilized metal affinity chromatography via the 8xHis tag. After elution, proteins were passed over a Superdex 200 Increase 10/300 GL (GE Healthcare) size exclusion column in PBS (Corning). Prior to immunization, 8xHis tags were cleaved using HRV 3C protease (ThermoScientific). Cleaved protein was repurified using Cobalt-TALON resin in order to remove the protease, cleaved tag, and any uncleaved protein. Approximate purified homotrimer yields per liter of Expi293F transfected were as follows: SARS-CoV-2 – 3 mg; SARS-CoV – 20 mg; WIV1-CoV – 25 mg.

### Fab and IgG Expression and Purification

Genes for the variable domains of the heavy and light chains were codon optimized by Integrated DNA Technologies and cloned into pVRC constructs containing the respective constant domains as previously described (62, 63). Heavy-chain Fab constructs contained a HRV 3C-cleavable 8xHis tag, while heavy-chain IgG constructs contained HRV 3C-cleavable 8xHis and SBP tags. Transfections and purifications were performed according to the same protocols used for the RBDs and homotrimers.

### Biolayer Interferometry

Biolayer interferometry (BLI) experiments were performed using a BLItz instrument (ForteBio). Fabs were immobilized on a FAB2G biosensor (ForteBio), and CoV proteins were the analytes. All proteins were diluted in PBS at room temperature. Titrations were performed to determine binding affinities. Single-hit concentrations at 10 μM were performed to get an approximate *KD*, and then subsequent titrations at appropriate concentrations (at least three). A final *KD* estimate was determined using a global fit model with a 1:1 binding isotherm using vendor-supplied software.

### Immunizations

C57BL/6 mice (Jackson Laboratory) received 20 μg of protein adjuvanted with 50% w/v Sigma Adjuvant System in 100 μL of inoculum (25). All immunizations were administered through the intraperitoneal route. Mice were primed (day 0) and received boosting immunizations at day 21 and day 42. Serum samples were collected on day 56 for characterization, with flow cytometry occurring between days 56 and 63. In this study, female mice aged 6-10 weeks were used. All experiments were conducted with institutional IACUC approval (MGH protocol 2014N000252).

### Flow Cytometry

Spleens were isolated from mice and single cell suspensions were generated by straining through a 70 μm cell strainer. Red blood cells were removed by treating with ACK lysis buffer and washed with PBS. Single cell suspensions were first stained with Aqua Live/Dead amine-reactive dye (0.025 mg/mL) before applying the following B and T cell staining panel using the staining approach described previously (25, 64). This included the following mouse-specific antibodies: CD3-BV786 (BioLegend), CD19-BV421 (BioLegend), IgM-BV605 (BioLegend), IgG-PerCP/Cy5.5 (BioLegend).

Streptavidin-conjugated fluorophores were used to label the SBP-tagged proteins as probes for flow cytometry. For the cohorts that received the SARS-CoV-2 spike prime followed by either the SARS-CoV-2 RBD homotrimer boost or the RBD homotrimer cocktail boost, the following probes were generated: SARS-CoV-2 RBD-APC/Cy7 (streptavidin-APC/Cy7 from BioLegend), SARS-CoV-2 spike-StreptTactin PE (StrepTactin PE from IBA Lifesciences), SARS-CoV-2 ΔRBM RBD-PE/Cy5.5 (streptavidin-PE/Cy5.5 from BioLegend). The panel for the cohort that received the SARS-CoV-2 spike prime followed by the RBD homotrimer cocktail boost also included SARS-CoV RBD-APC (streptavidin-APC from BioLegend) and WIV1-CoV RBD-BV650 (streptavidin-BV650 from BioLegend). For the cohort that received three SARS-CoV-2 spike immunizations, the following probes were generated: SARS-CoV-2 RBD-APC/Cy7 (streptavidin-APC/Cy7 from BioLegend), SARS-CoV-2 spike-StreptTactin PE (StrepTactin PE from IBA Lifesciences), SARS-CoV-2 ΔRBM RBD-APC (streptavidin-APC from BioLegend). Conjugations were performed as previously described (65). Briefly, fluorescent streptavidin conjugates were added in 5 increments with 20 minutes of incubation with rotation at 4°C in between to achieve a final molar ratio of probe to streptavidin valency of 1:1. The final conjugated probe concentration was 0.1 μg/mL. Flow cytometry was performed on a BD FACSAria Fusion cytometer (BD Biosciences). Analysis of the resultant FCS files was conducted using FlowJo (version 10).

### Serum ELISAs

Serum ELISAs were performed by coating Corning 96-well clear flat bottom high bind microplates with 100 μL of protein at 5 μg/mL in PBS. Plates were incubated overnight at 4°C. Coating solution was removed, and plates were blocked using 1% BSA in PBS with 1% Tween for 60 minutes at room temperature. Blocking solution was removed. Sera were diluted 1:40 in PBS, and 5-fold serial dilution was performed. CR3022 IgG at a starting dilution of 5 μg/mL with 5-fold serial dilution was used as a positive control. 40 μL of primary antibody solution was applied to each well. Primary incubation occurred for 90 minutes at room temperature. Plates were then washed three times with PBS-Tween. HRP-conjugated rabbit anti-mouse IgG antibody (Abcam) at a concentration of 1:20,000 in PBS and a volume of 150 μL was used as a secondary antibody. Secondary incubation occurred for 60 minutes at room temperature. Plates were then washed three times with PBS-Tween. 1xABTS development solution (ThermoFisher) was applied as outlined in the manufacturer’s recommendations. Development was stopped after 30 minutes with a 1% SDS solution. Plates were read at 405 nm using a SectraMax iD3 plate reader (Molecular Devices).

### Competition ELISAs

Competition ELISAs were performed using a similar protocol to serum ELISAs. The primary incubation consisted of 40 μL of the relevant IgG at 1 μM. Incubation occurred at room temperature for 60 minutes. Mouse sera were then spiked in at a final concentration in the linear range of the serum ELISA titration curve (1:800 for the cohort that received three SARS-CoV-2 RBD monomer immunizations, 1:12,800 for all other cohorts). Plates were incubated at room temperature for an additional 60 minutes. The primary solution was removed, and plates were washed three times using PBS-Tween. HRP-conjugated goat anti-mouse IgG, human/bovine/horse SP ads antibody (Southern Biotech) was applied at a concentration of 1:4000 and a volume of 150 μL as a secondary antibody. Plates were then incubated, washed, and developed using the same procedure as the serum ELISAs.

### ACE2 Cell Binding Assay

ACE2 expressing 293T cells (66) (a kind gift from Nir Hacohen and Michael Farzan) were harvested and washed with PBSF. Cells were allocated such that 200,000 cells were labelled for each condition. Cells were incubated with 100 μL of 200 nM antigen in PBS for 60 minutes on ice. Following two washes with PBS supplemented with 2% FBS, cells were incubated with 50 μL of 1:200 streptavidin-PE (Invitrogen) on ice for 30 minutes. Cells were washed twice and resuspended in 100 μL of PBS supplemented with 2% FBS. Flow cytometry was performed using a Stratedigm S1000Exi Flow Cytometer. FCS files were analyzed using FlowJo (version 10).

### Human Serum Samples

A total of 24 human serum samples were obtained from a previously characterized cohort (9). All patients had received either 1 or 2 doses of the Pfizer (BNT162b2) or Moderna (mRNA-1273) vaccines. Of the patients who had received both doses, samples were collected a median of 14 days following the second dose (range: 4 days – 32 days). The median patient age was 51 years, with ages ranging from 22-66 years.

### Serum Adsorption

SNAP-tagged (61) MERS-CoV RBD, SARS-CoV-2 ΔRBM RBD with four additional putative glycosylation sites (Figure S5), and SARS-CoV-2 RBD were conjugated to SNAP-Capture Pull-Down resin (New England BioLabs). For each conjugation, 20 μL of settled resin was resuspended in 100 μL of protein at 1 mg/mL per the manufacturer’s recommendations. Both MERS-CoV RBD and SARS-CoV-2 RBD conjugation reactions also included 1 mM DTT to improve conjugation efficiency. Wildtype-like reactivity to conformationally specific Fabs (CR3022, S309, and B38 for SARS-CoV-2 RBD; m336 for MERS RBD) in the presence of 1 mM DTT was confirmed prior to conjugation (15, 17, 18, 67). Conjugation occurred with rotation overnight at 4°C.

Following conjugation, the resin was washed 5 times with PBS via resuspension followed by centrifugation and the removal of the supernatant. Sera from the mice in each cohort were pooled and diluted 1:40 in PBS to a total volume of 100 μL. Sera from human patients were diluted 1:20 in PBS to a total volume of 100 μL. Diluted sera were added to the conjugated resin and incubated with rotation overnight at 4°C. Resin was filtered from the sera following incubation, and the adsorbed sera was used for neutralization assays and ELISAs.

### Serum Adsorption ELISAs

Serum adsorption ELISAs were performed using a similar protocol to the serum ELISAs. For the primary antibody incubation, adsorbed serum solution was diluted 1:2, and subsequent serial 5-fold dilutions were generated. The remainder of the assay was performed according to the serum ELISA procedure.

### Pseudovirus Neutralization Assay

Serum neutralization against SARS-CoV-2, SARS-CoV, and WIV1-CoV was assessed using lentiviral particles pseudotyped with the respective spike proteins as previously described (8, 9). Lentiviral particles were produced via transient transfection of 293T cells. The titers of viral supernatants were determined via flow cytometry on 293T-ACE2 cells (66) and via the HIV-1 p24^CA^ antigen capture assay (Leidos Biomedical Research, Inc.). Assays were performed in 384-well plates (Grenier) using a Fluent Automated Workstation (Tecan). For mouse sera, samples were initially diluted 1:9, with subsequent serial 3-fold dilutions. For human and mouse sera that had previously been adsorbed, the adsorbed sample was used at an initial 1:3 dilution, and serial 3-fold dilutions were performed. Serum sample volume in each well was 20 μL, and 20 μL of pseudovirus containing 125 infectious units was added. The combination was incubated for 60 minutes at room temperature. Afterwards, 10,000 293T-ACE2 cells (66) in 20 μL of media containing 15 μg/mL polybrene was added. The plates were then incubated at 37°C for 60-72 hours.

A previously described assay buffer was used to lyse the cells (68). A Spectramax L luminometer (Molecular Devices) was used to quantify luciferase expression. Percent neutralization at each serum concentration was determined by subtracting background luminescence from cells only sample wells, then dividing by luminescence of wells with only virus and cells. GraphPad Prism was used to fit nonlinear regressions to the data, which allowed IC50 values to be calculated using the interpolated 50% inhibitory concentration. IC50 values were calculated for all samples with a neutralization value of at least 80% at the highest serum concentration.

### Phylogenetic Trees

Prior to generating phylogenetic trees, alignments were generated via ClustalOmega. Phylogenetic trees were then generated using those alignments as inputs into a neighbor-joining algorithm via ClustalW2. All settings were left as default.

### Statistical Analysis

Statistical analyses and curve fitting were performed using GraphPad Prism (version 9). To compare two populations of continuous variables without evidence of conforming to a normal distribution, the non-parametric two-tailed Mann-Whitney U test was used. To compare multiple populations meeting this description, the Kruskal-Wallis test was used with post hoc analysis using Dunn’s multiple comparison testing. The ratio paired t-test was used to compare two populations with evidence of normality. P values were corrected for multiple comparisons, and a p value < 0.05 was considered significant.

## Acknowledgments

We thank members of the Schmidt and Lingwood Laboratories for helpful discussions. We thank Dr. Jason McLellan from University of Texas, Austin for the spike plasmid. We thank Nir Hacohen and Michael Farzan for the kind gift of the ACE2 expressing 293T cells to ABB.

## Author contributions

Conceptualization, BMH, MS, JF, TMC, DL, AGS; Methodology, BMH, ECL, TMC, ABB, DL, AGS; Investigation, BMH, MS, ECL, KSD, JF, ASY, TK; Writing – Original Draft, BMH and AGS; Writing – Review and Editing, all authors; Funding Acquisition, ABB, DL, AGS; Supervision, ABB, DL, AGS.

## Data and materials availability

All data is available in the main text or in the supplementary materials.

