## Supplement for "Engineered receptor binding domain immunogens elicit pan-sarbecovirus neutralizing antibodies outside the receptor binding motif"

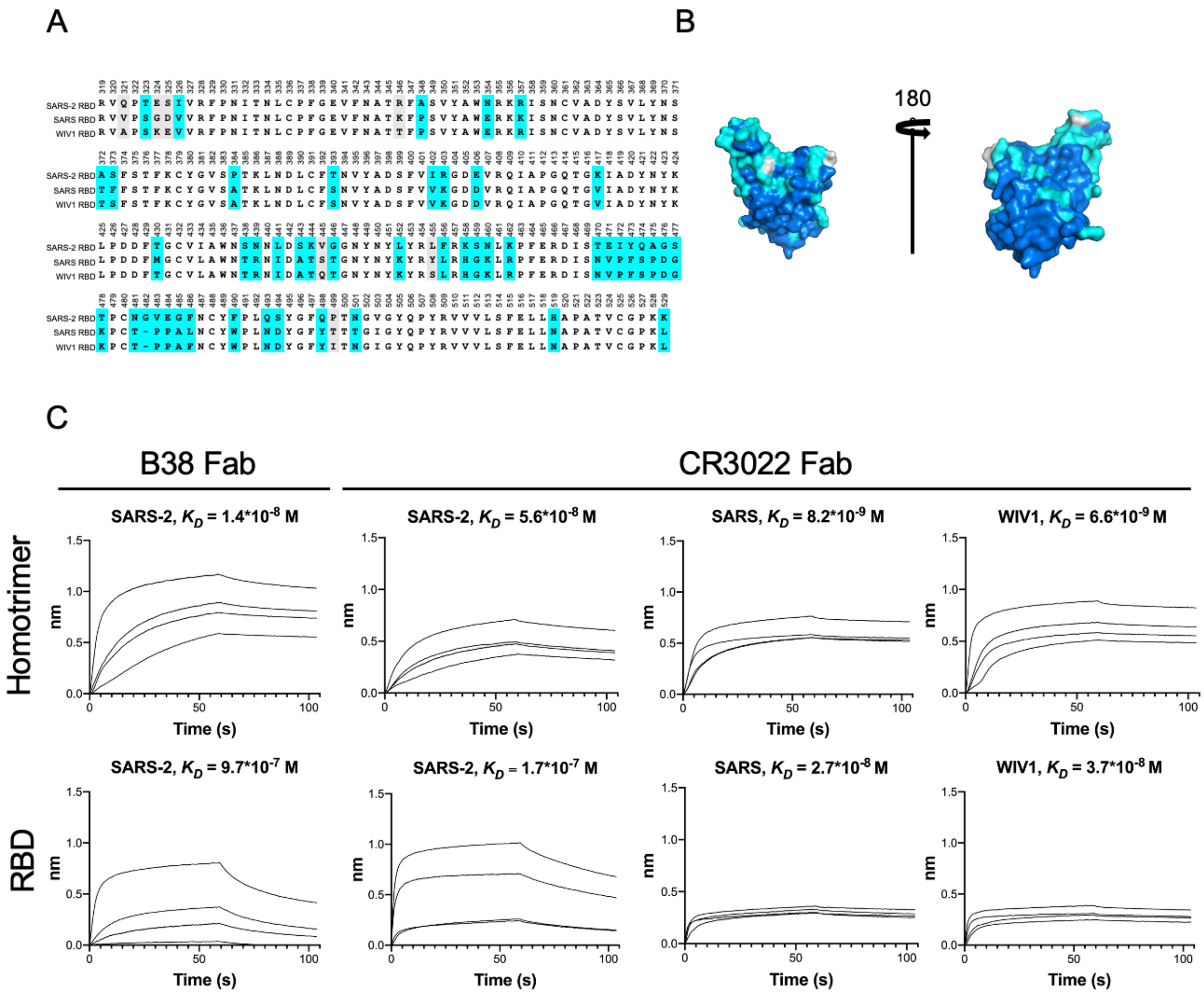

**Fig. S1. Sequence Conservation and BLI with conformationally-specific Fabs.** (**A, B**) Strict amino acid conservation across the SARS-CoV-2 RBD (Genbank MN975262.1), SARS-CoV RBD (Genbank ABD72970.1), and WIV1 RBD (Genbank AGZ48828.1) is depicted using dark blue on the structure and white in the table for matches between all three genes, light blue for matches between two genes, and silver for positions where all genes differ (PDB: 6M0J). (**C**) Conformationally specific Fabs CR3022 and/or B38 were used to verify that homotrimer affinity was comparable (or greater than, due to increased avidity) wildtype RBD affinity. Fabs were immobilized to FAB2G sensors, and coronavirus proteins were the analytes. Homotrimers were titrated at 1 μM, 750 nM, 500 nM, and 250 nM. Monomeric SARS-CoV and WIV1-CoV RBDs were titrated at 10 μM , 5 μM, 2.5 μM, and 1 μM. Monomeric SARS-CoV-2 RBD was titrated at 10 μM, 1 μM, 500 nM, and 100 nM with B38 Fab and at 10 μM, 5 μM, 750 nM, and 250 nM with CR3022 Fab. Apparent $K_D$ was obtained by vendor-supplied software. (Related to **Fig. 1**)

A

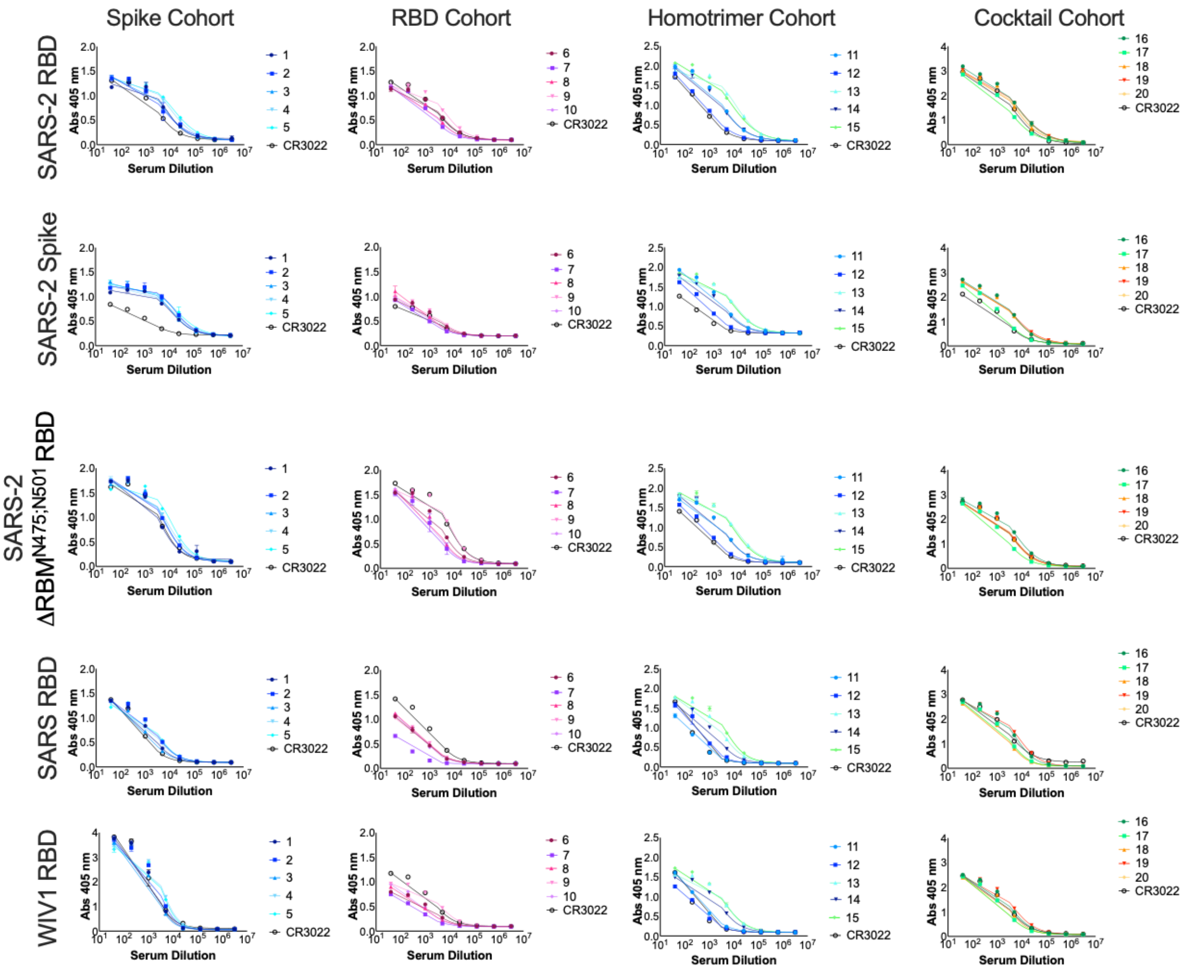
Spike Cohort
RBD Cohort
Homotrimer Cohort
Cocktail Cohort
SARS-2 RBD
SARS-2 Spike
SARS-2 ΔRBM N475;N501 RBD
SARS RBD
WIV1 RBD
Abs 405 nm
Serum Dilution
CR3022

B

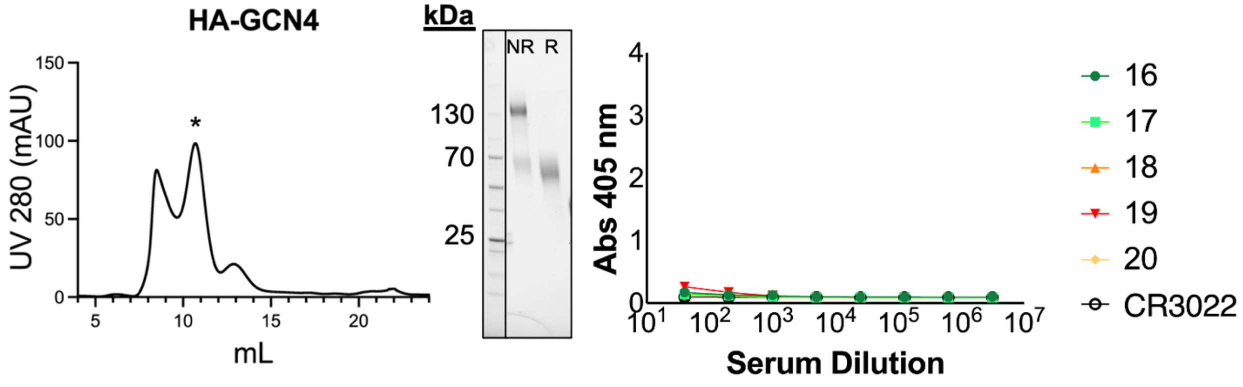
HA-GCN4
UV 280 (mAU)
mL
kDa
NR
R
130
70
25
Abs 405 nm
Serum Dilution
16
17
18
19
20
CR3022

C

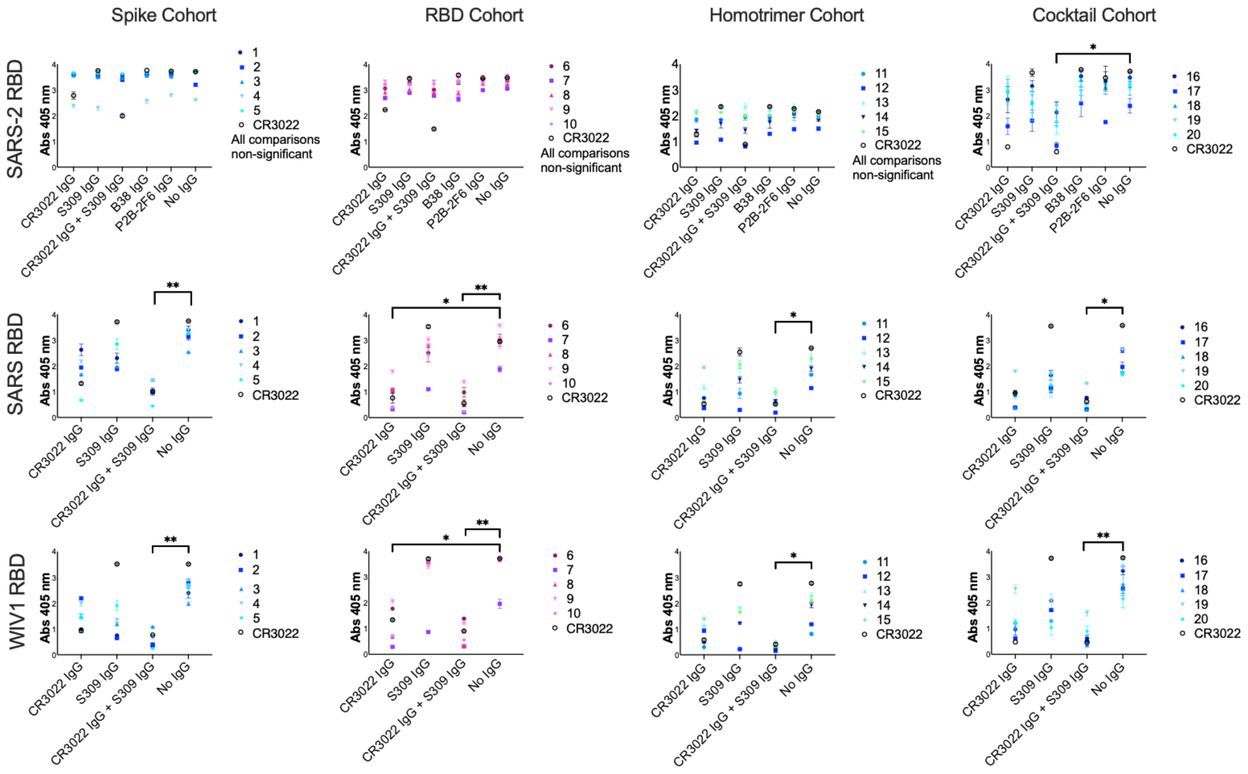
Spike Cohort
RBD Cohort
Homotrimer Cohort
Cocktail Cohort
SARS-2 RBD
SARS RBD
WIV1 RBD
Abs 405 nm
CR3022 IgG
S309 IgG
CR3022 IgG + S309 IgG
B38 IgG
P2B-2F6 IgG
No IgG
1
2
3
4
5
CR3022
All comparisons non-significant
*
**

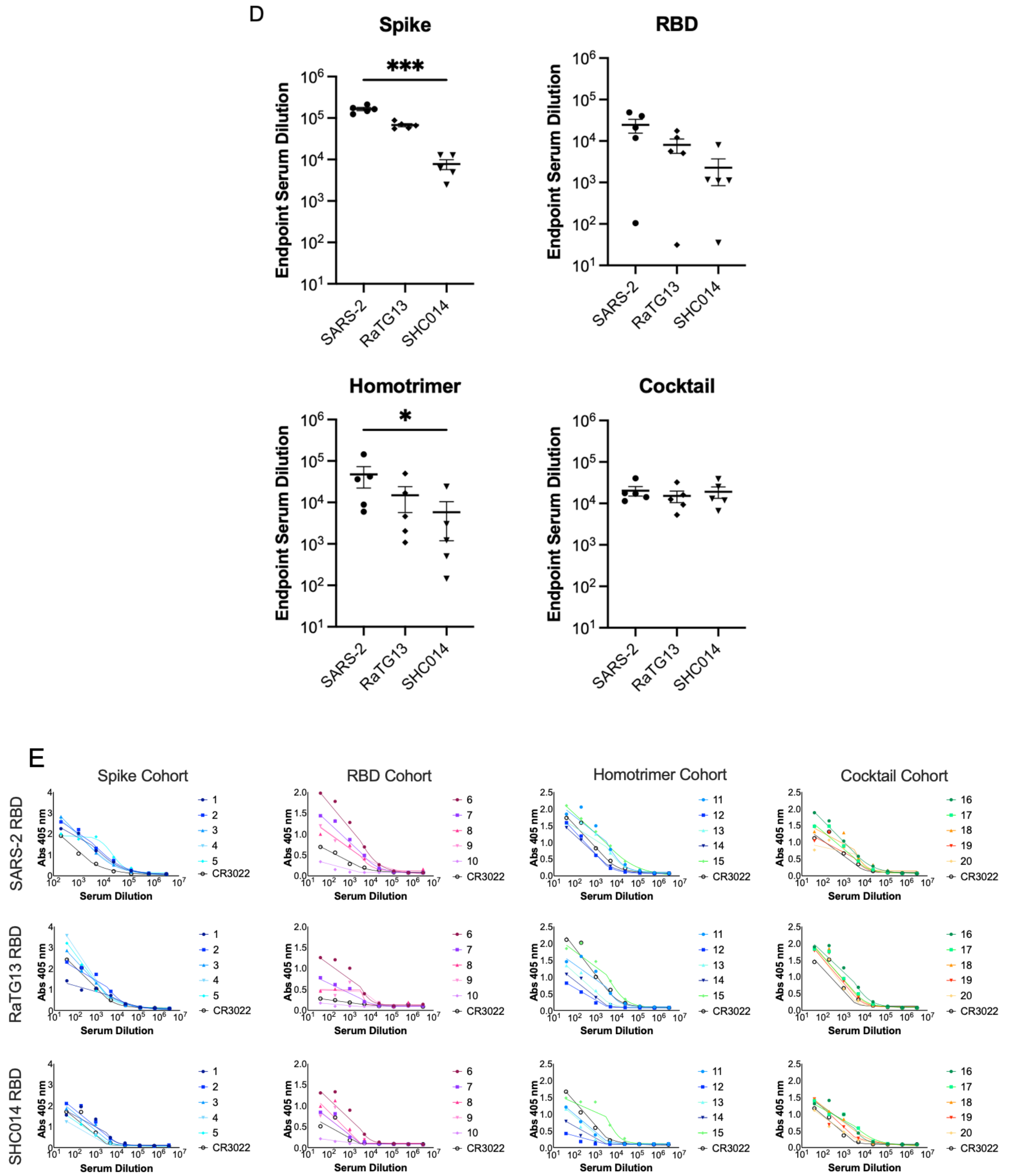

**Fig. S2. Serum ELISAs.** (**A**) Serum ELISAs for each cohort against different coronavirus proteins as coating antigens, with endpoint titers shown in **Figure 2B-C.** Curves were fit using a sigmoidal model and are shown. **(B)** A serum ELISA was performed for the cohort that received the RBD homotrimer cocktail boost with an irrelevant protein (influenza hemagglutinin head) tagged with the hyperglycosylated GCN4 tag (HA-GCN4) to measure tag-directed antibody responses. For the coating HA-GCN4 protein, a purification size exclusion trace (fractions in the peak marked with "*" were pooled) and an SDS-PAGE gel run under non-reducing (NR) and reducing (R) conditions are shown. (**C**) Competition ELISAs against a panel of SARS-CoV-2, SARS-CoV, and WIV1-CoV-directed IgGs CR3022, S309, with full results shown here corresponding to **Figure 2D**. (**D**) Serum was assayed in ELISA at day 56 with different coronavirus RBDs. Statistical significance was determined using Kruskal-Wallis test with post-hoc analysis using Dunn's test corrected for multiple comparisons ( * = $p < 0.05$, ** = $p < 0.01$, *** = $p < 0.001$). (**E**) Curves corresponding to endpoint titers in Fig. S2D. Curves were fit using a sigmoidal model. (Related to **Fig. 2**)

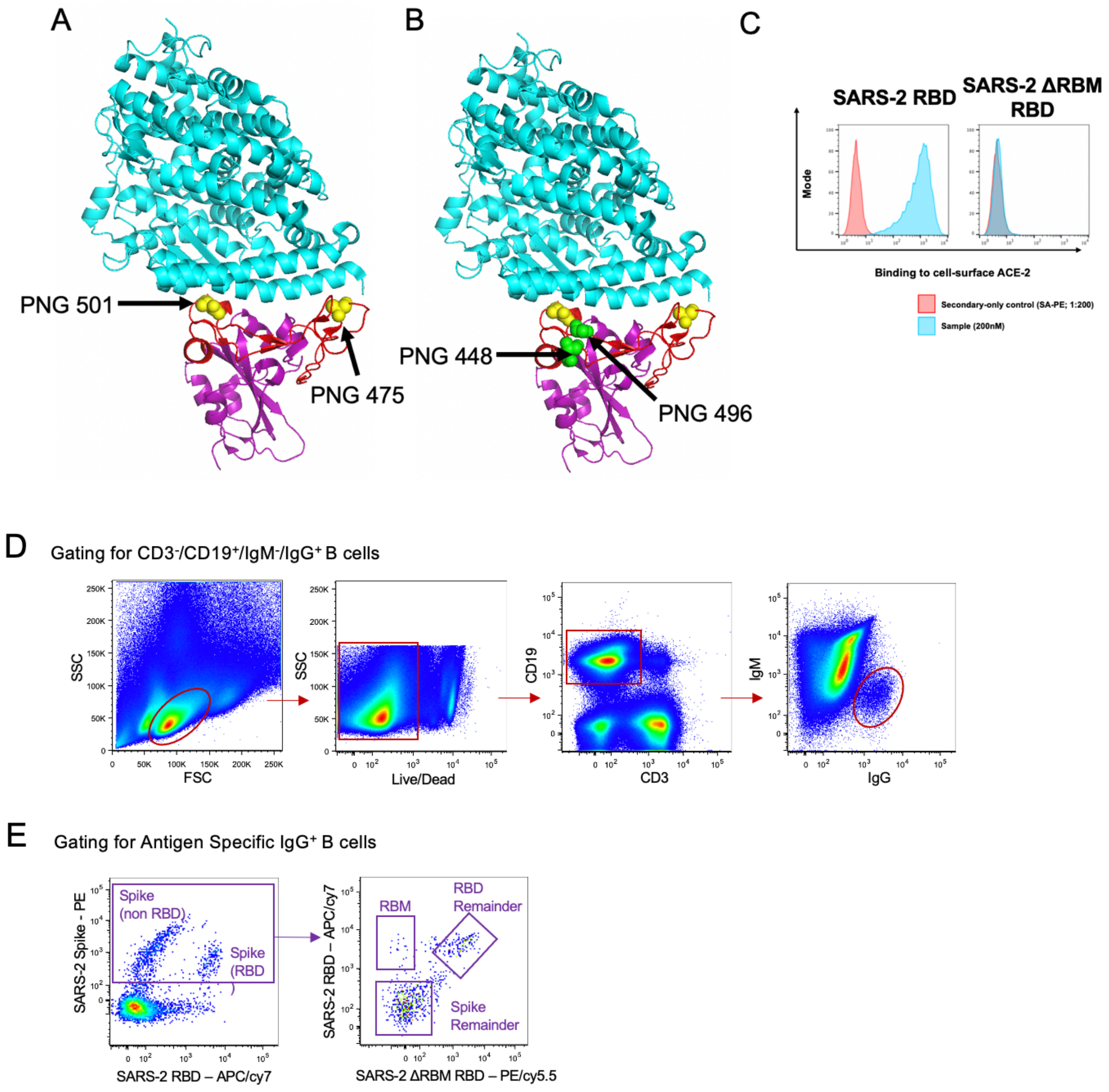

**Fig. S3. Validation of SARS-CoV-2 ΔRBM constructs and flow cytometry gating scheme.** (**A, B**) Design and expression of two different versions of the SARS-CoV-2 RBD with additional putative N-linked glycosylation sites (PNGs) engineered onto the RBM. One construct has glycosylation sites at positions 475 and 501, while the other construct has glycosylation sites at positions 475, 501, 448, and 496. (PDB: 6M0J) (**C**) These constructs were both termed SARS-

CoV-2 ΔRBM RBD to indicate the loss of ACE2 binding to the RBM. (**D**) Gating strategy to select for CD3-/CD19+/IgM-/IgG+ cells in order to isolate the memory B cell population for further analysis. (**E**) Antigen specific memory IgG responses were identified using a combination of SARS-CoV-2 spike, SARS-CoV-2 RBD, and SARS-CoV-2 ΔRBM RBD flow hooks. SARS-CoV-2 spike-directed B cell responses were further separated by reactivity to into the RBM, RBD remainder (excluding RBM epitopes), and spike remainder (excluding RBM and RBD epitopes). (Related to **Fig. 3**)

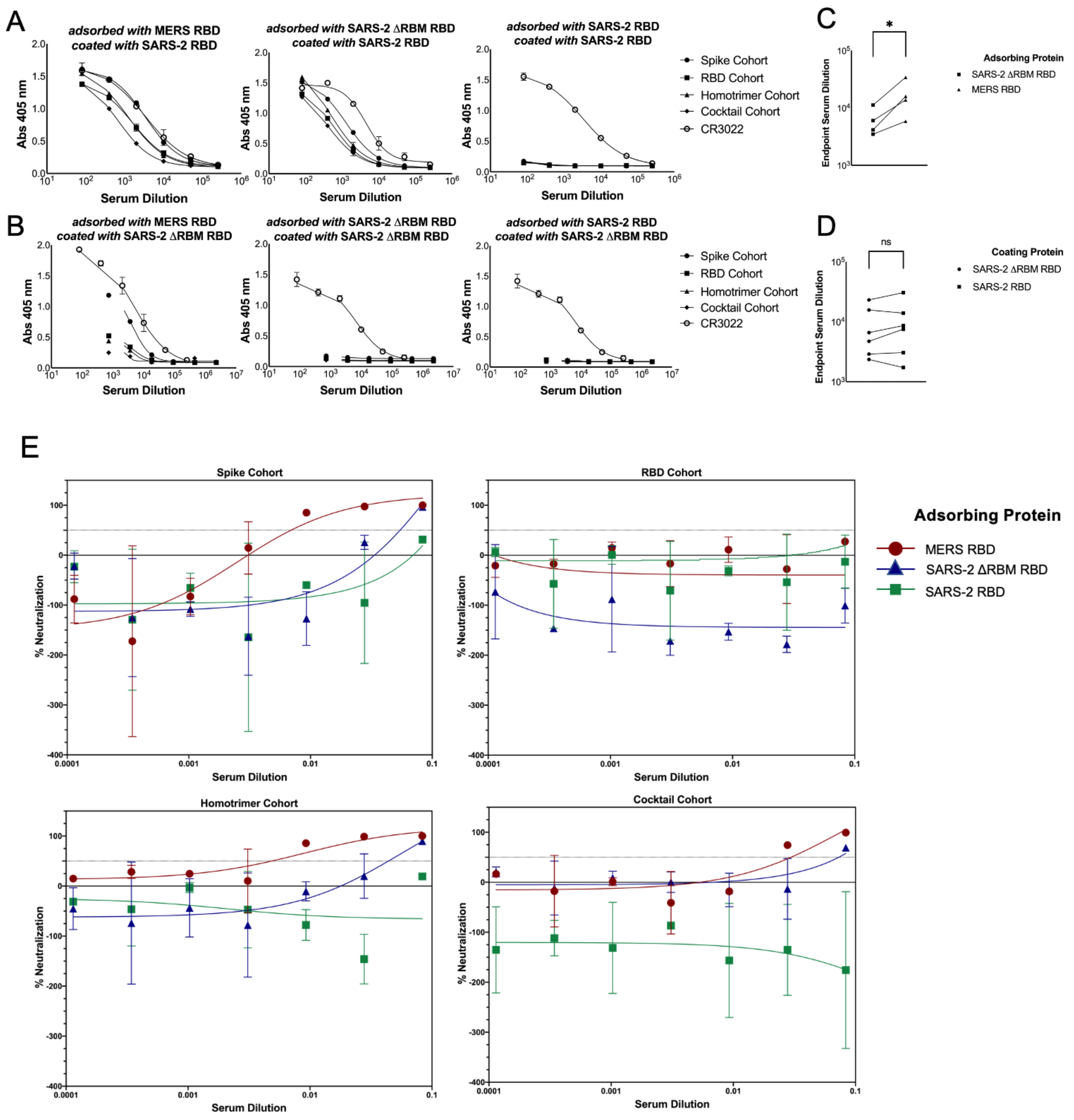

**Fig. S4. Serum adsorption validation and neutralization results.** (**A, B**) Data shows two rounds of adsorption of pooled sera from each cohort with MERS-CoV RBD, SARS-CoV-2 ΔRBM RBD, or SARS-CoV-2 RBD. ELISAs were performed with the adsorbed sera using SARS-CoV-2 RBD and SARS-CoV-2 ΔRBM RBD as coating antigens. (**C**) Sera adsorbed with the SARS-CoV-2

ΔRBM RBD lost a significant proportion of their SARS-CoV-2 RBD-directed antibodies compared to the negative control adsorbed with MERS-CoV RBD (ratio paired t-test, $p = 0.0129$) and nearly all of their non-RBM directed antibodies. (**D**) Sera adsorbed with MERS-CoV RBD did not show a significant difference in endpoint titers against SARS-CoV-2 ΔRBM RBD and SARS-CoV-2 RBD. **(E)** Raw neutralization assay data and nonlinear curve fits from pooled serum samples following adsorption with MERS-CoV RBD, SARS-CoV-2 ΔRBM RBD, or SARS-CoV-2 RBD. The blue curve in the top left panel serves as an example of a curve that showed some neutralization activity but for which an NT50 value could not be fit. (Related to **Fig. 4**)
